# Field evaluation of biocontrol agents against black-foot and Petri diseases of grapevine

**DOI:** 10.1101/2020.05.19.101568

**Authors:** María del Pilar Martínez-Diz, Emilia Díaz-Losada, Marcos Andrés-Sodupe, Rebeca Bujanda, María Mercedes Maldonado-González, Sonia Ojeda, Amira Yacoub, Patrice Rey, David Gramaje

## Abstract

**BACKGROUND:** Black-foot and Petri diseases are the main fungal diseases associated with young grapevine decline. Two field experiments were established to evaluate the preventive effect of two potential biocontrol agents (BCAs), i.e. *Streptomyces* sp. E1 + R4 and *Pythium oligandrum* Po37, and three BCA-commercial products containing *Trichoderma atroviride* SC1, *Trichoderma koningii* TK7 and *Pseudomonas fluorescens*+*Bacillus atrophaeus* on fungal infection in grafted plants and plant growth parameters.

**RESULTS:** The effectiveness of some BCA in reducing the incidence and severity of both diseases was dependent on the plant part analyzed and the plant age. No single BCA application was able to control both diseases. *Streptomyces* sp. E1+R4 were able to reduce significantly black-foot disease infection while *P. oligandrum* Po37 and *Trichoderma* spp. were able to reduce significantly Petri disease infection. BCA treatments had no effect on the shoot weight, and root weight was significantly lower in all BCA treatments with respect to the control.

**CONCLUSIONS:** The combination of the disease-suppressive activity of two or more beneficial microbes in a biocontrol preparation is required to prevent infection by black-foot and Petri disease fungi in vineyards.

## 1 INTRODUCTION

Grapevine trunk diseases (GTDs) are one of the most damaging diseases affecting the grapevine industry in all grape-growing regions worldwide, being responsible for yield and productivity loss, and one of the main causes of an early vines death.^1^ Among them, black-foot and Petri diseases are the two most common GTDs affecting planting material at nurseries, newly planted vines and young vineyards (<5 years old).^2-4^ In La Rioja (northern Spain), the annual financial cost of the replacement of death plants cv Tempranillo in due to black-foot and Petri diseases is estimated to be 7.16 million €/year.^5^ Field symptoms of black-foot and Petri diseases affected vines include overall stunting growth, delayed budbreak, retarded or absent sprouting, shortened internodes, chlorotic and sparse foliage with necrotic margins, leaves or entire shoots wilting, and dieback.^3^ However, these symptoms also resemble those associated with abiotic disorders such as spring frost, winter damage and/or nutrient deficiency.^1^ Characteristic symptoms of black-foot disease include dark brown and soft areas in roots and black discolouration and necrosis in the basal end of the rootstock.^2^ Regarding Petri disease, dissected affected vines display brown and black vascular streaking, mainly in the rootstock, and gumming that turns dark when exposed to air.^3^

Up to 32 species of the genera *Campylocarpon, Cylindrocladiella, Dactylonectria, Ilyonectria, Neonectria, Pleiocarpon* and *Thelonectria* have been reported to cause black-foot disease,^1,6-8^ *Dactyonectria torresensis* being the most prevalent species associated with diseased vines in Europe.^9-11^ These fungal species are known to be soilborne and persist as mycelium and conidia in rotten root fragments or as resting spores (chlamydospores) that can survive in the soil for extended periods of time after infected plants are removed.^4,12^ Apparently healthy plants placed in infested nursery soil can become infected through trunk wounds or roots, such as the incomplete callused rootstock end.^2^

The main fungal species associated with Petri disease is *Phaeomoniella (Pa.) chlamydospora*.^13^ However, other fungal species that have also been isolated in relatively high frequencies from Petri diseased vines are 29 species of the genus *Phaeoacremonium, Pleurostoma richardsiae*, and 6 *Cadophora* spp.^1^ Among those, *Phaeoacremonium (Pm.) minimum* and *Cadophora luteo-olivacea* are the most prevalent.^13,14^ *Pa. chlamydospora* and *Phaeoacremonium* spp. can spend part of their disease cycle in soil as mycelium and conidia in infected rootstock wood, roots or pruning debris,^15,16^ or chlamydospores in the case of *Pa. chlamydospora*.^15^ The presence of *Cadophora* spp. in soils has been recently confirmed using ITS high-throughput amplicon sequencing (HTAS) approach.^17^ Therefore, the main hypothesis is that these fungi could gain entry into the xylem of young plants at the nursery or newly established vineyards through root and/or basal end of the rootstock infections. In addition, they are also disseminated through the dispersion of airborne spores (conidia and/or ascospores) by rain, wind or arthropods until they land on susceptible and fresh pruning wounds.^15^

Presently, no curative measures are available to reduce the impact of these diseases once the vines are infected, making their management in the field difficult. Furthermore, the loss of the most effective preventative chemical products such as the banning in the early 2000s of sodium arsenite or benzimidazoles,^18^ and the high current restrictions and difficulties that chemicals are facing in most countries around the world because of the risks for human health and the environment,^19,20^ increase even more the complexity of their control. Nowadays, the best way to handle these diseases is by using an Integrated Pest Management (IPM) strategy^21^ where several strategies are combined to reduce GTDs infections, such as the use of physical (e.g. hot-water treatment), biological (e.g. antagonist microorganisms) and cultural practices (e.g. crop management, irrigation, soil preparation, etc.), throughout the nursery mother blocks and newly planted vineyards.^1^

Investigation of BCAs able to prevent or at least reduce the development of GTDs are considered a research priority.^1^ In fact, over the last 10 years there has been a frantic search by the GTDs research community for microbial antagonists, including fungi,^21-33^ bacteria,^29,34-40^ and oomycetes.^41,42^ Although some of these studies provided promising findings, the results have not been consistent, observing differences in efficacy depending on the nature of the BCA, the target pathogen, application method, time of exposure to the BCA and even the grapevine cultivars and rootstocks subjected to study. In addition, most of these studies have been performed so far under *in vitro* laboratory,^26,29,31,32,34-40^ greenhouse^29,31,32,35-37,41,42^ or nursery^21-25,28,33,38^ controlled conditions by using rootstock or scion cuttings.

Three *Trichoderma*-based biological products are currently registered in Spain for the preventive protection of pruning wounds against GTD fungi, namely Esquive® (*Trichoderma atroviride* I-1237), Blindar® (*Trichoderma asperellum* ICC012 + *Trichoderma gamsii* ICC080) and Vintec® (*T. atroviride* SC1).^43^ Only Vintec® has been additionally registered to control Petri disease pathogens in grapevine grafted nursery stock.^43^ Therefore, we propose to apply registered BCA products in Spain for control of GTD fungi both on grapevine and/or other hosts, and other potential BCAs as a preventive strategy in pre- and post-planting. The main objectives of this study were: (i) to evaluate the effectiveness of several BCA root treatments under field conditions in reducing natural infections of fungal pathogens associated with black-foot and Petri diseases over two growing seasons, and (ii) to assess the BCA root treatments influence in plant growth parameters.

## 2 MATERIALS AND METHODS

### 2.1. Planting material

One-year old grapevine grafted plants of ‘Tempranillo’/110 Richter combination with uniform root distribution were obtained from a commercial nursery in Spain and used in this experiment. Roots were trimmed to 10 cm length and dormant plants were hot-water treated at 53°C for 30 min to reduce any existing infections by black-foot and Petri disease pathogens^44,45^ and then acclimatized for 24 h at 20°C before biological control agents (BCA) inoculation.

### 2.2. Grafted plants inoculation and experimental design

Hot-water treated plants were inoculated by dipping the roots and the basal part of the plants for 24 h at room temperature with 25 l water suspensions of the following treatments: (T1) *Streptomyces* sp. E1 + R4 (1.35 x 10^9^ CFU ml^-1^) at 7.5 ml l^-1^, (T2) *Trichoderma koningii* TK7 (Condor Shield®, ATENS; 1 x 10^9^ CFU g^-1^ formulated product) at 2 g l^-1^, (T3) *T. atroviride* SC1 (Vintec®, Belchim Crop Protection; 2 x 10^10^ CFU g^-1^ formulated product) at 2 g l^-1^, (T4) *Pseudomonas fluorescens + Bacillus atrophaeus* (Stilo Cruzial®, SIPCAM Iberia; 1x 10^8^ CFU g^-1^ formulated product) at 2 g l^-1^, (T5), *Pythium oligandrum* Po37 (Biovitis, France; 1.28 x 10^6^ CFU g^-1^) at 2 g l^-1^, and (C) water as untreated control. We selected T1 and T5 due to the previously demonstrated efficacy against GTD fungi in young vines.^38,41^ T2 and T4 are not registered as a phytosanitary product in Spain yet. The viability of the *Trichoderma* conidia in the products T2 and T3 was checked to be at a minimum of 85% before the trial, as described by Pertot *et al*.^28^

Inoculated grafted plants were immediately planted in May 2017 in two field sites located in Logroño (La Rioja, Spain). Both fields were under grapevine nursery planting material rotation, which is very common in the area of study. Standard cultural practices were used in both sites during the grapevine growing season. The plant groups (40 plants) were spaced 100 cm from other groups, plants being 30 cm apart from center to center. Each field plot was 12 m long and included 24 rows, each with a plant group of 40 plants (960 plants per field). In both sites, the experimental design consisted of four randomized blocks, each containing a plant group (40 plants) of each treatment (160 plants per treatment), with 200 cm between each block. Plots were less than 1 km apart and had very similar climates. Soil samples were taken for physicochemical properties analysis as described below. A drip irrigation system was laid on the soil of each row. An additional stock of 50 grafted plants was used to check for their phytosanitary status immediately after hot-water treatment (HWT).

### 2.3. GTD fungal isolation and identification

In February 2018, once grafted plants had completed their cycle of vegetative growth and were in a dormant state, 50% of the 2-year-old plants in each field were carefully dug out from the soil to keep the root system intact and taken back to the laboratory for immediate processing. In order to isolate black-foot and Petri disease pathogens, two plant parts were evaluated, roots and the basal ends of the rootstocks. Root necrotic sections from 2-3 cm near the basal end of the rootstock and wood sections of 3 cm length of the basal end of the rootstock were cut, washed under running tap water, surface sterilized in 33% sodium hypochlorite (commercial 40 g Cl/l) for 1 min and rinsed twice with sterile distilled water. Five small root or xylem pieces were plated on Malt Extract Agar (MEA) supplemented with 0.35 gl^−1^ of streptomycin sulphate (Sigma-Aldrich, St. Louis, MO, USA) (MEAS). Four MEAS plates were used per plant (two per plant part). Plates were incubated for 10-15 days at 25 °C in the dark and all colonies were transferred to Potato Dextrose Agar (PDA). Isolates were single-spored prior to morphological and molecular identification with the serial dilution method.^46^

In May 2018, the remaining 50% of the plants in each field were drip inoculated with all treatments (0.5 l per plant using the same inoculum concentration as described above). In February 2019, these 3-year-old plants were carefully dug out and processed for fungal isolation as described above. All planting material was washed and also assessed for undried shoot and total root weight. The disease incidence (DI) of black-foot and Petri disease pathogens was determined as the mean percentage of grafted plants that was infected by these fungi. The disease severity (DS) in infected grafted plants was determined as the mean percentage of root or wood segments (ten segments per plant each) that was colonized by these fungi. The presence of *Trichoderma* spp. was also recorded to provide an indication of the extent of colonization following treatment with the *Trichoderma* formulations (T2 and T3). The stock of 50 plants was also analysed after HWT as described before.

Fungal isolates resembling black-foot and Petri disease pathogens were identified by molecular techniques. For DNA extraction, 300 mg of fungal mycelium and conidia from single spore isolates grown on PDA for 2 to 3 weeks at 25°C in the dark were scraped and homogenised twice in a Fastprep®-24 tissue homogenizer (MP Biomedicals, USA). Total DNA was extracted using the E.Z.N.A. Plant Miniprep Kit (Omega Bio-tek, Doraville, USA) following manufacturer’s instructions. DNA was visualized on 1% agarose gels stained with RedSafe (iNtRON Biotechnology, Lynnwood, WA, USA). DNA was stored at −20°C. Black-foot species were identified by sequencing part of the histone gene (*his*3) using CYLH3F and CYLH3R primers.^47-49^ The identification of *Pa. chlamydospora* isolates was performed by analysis of the ITS region of DNA amplified using the fungal primers Pch1/Pch2.^50^ *Pm. minimum* and *C. luteo-olivacea* were identified by sequence analysis of the β-tubulin (*tub2)* using the primer pairs T1^51^ and Bt2b^52^ for *Phaeoacremonium*, and BTCadF/BTCadR^53^ for *Cadophora. Trichoderma* spp. were isolated on MEAS and identified at species level by sequencing the ITS region using the universal primers ITS1F/ITS4.^54^ *P. oligandrum* was isolated on Corn Meal Agar added with Pimaricin, Ampicilliun, Rifampicin and Pentachloronitrobenzene (CMA-PARP) and identified by morphological features.^41^ Polymerase chain reaction (PCR) products were purified with the High Pure PCR Product Purification Kit (Roche Diagnostics, Mannheim, Germany), and sequenced in both directions by Macrogen Inc. (Seoul, Republic of Korea). The sequences obtained were then blasted in GenBank.

### 2.4. Soil physicochemical properties analysis

Four soil cores were collected to a depth of 20 cm from each field and bulked into a single soil sample per field. Samples were mixed well, air-dried for one week and sieved (2-mm to 5-mm mesh size) prior to soil physicochemical analyses. Soil samples were tested for electric conductivity (EC) in water and pH with a soil solution ratio of 1:5, soil texture by laser diffraction particle size (Diffractometer LS 13 320, Beckman Coulter Inc., Brea, Calif.), soil organic matter (SOM) by dichromate oxidation,^55^ cation exchange capacity (CEC) by the cobaltihexamine method,^56^ carbonate total by infrared (Equilab CO-202; Equilab, Jakarta, Indonesia), assimilable magnesium and calcium by inductively coupled plasma (ICP) spectroscopy (ARL-Fison 3410, USA) and the cobaltihexamine method and P, K, S, Mg, Mn, Fe, Ca and Na by ICP and Mehlich method.^57^ Analyses were conducted in the official Regional Laboratory of La Grajera (Logroño, Spain) in April 2017, before the beginning of the experiment.

### 2.5. Data analysis

Prior to statistical analyses, data were checked for normality and homogeneity of variances, and transformed when needed. Percentage data were transformed into arcsin (DI or DS/100)^1/2^. Each treatment means (DI, DS, root and shoot weights) was calculated from the corresponding values in each sampling moment. The statistical analysis of the experimental results was carried out in a two-way ANOVA with blocks and treatments as independent variables, and the following dependent variables: DI (%), DS (%), root weight (g) and shoot weight (g). In the 3-year-old plants, the percentage of reduction (PR) of the fungal pathogen detection at each isolation plant part and for each fungal GTD species was calculated as PR = 100(PC – PT)/PC, where PC is the mean pathogen incidence or severity in the control and PT is the mean pathogen incidence or severity in the BCA treatment. Means were compared by the Student’s *t* least significant difference (LSD) at *P* < 0.05. Soil physicochemical variables were subjected to analyses of variance. LSD test was calculated to compare variable means. Data from all experiments were analysed using the Statistix 10 software (Analytical Software).

## 3 RESULTS

### 3.1 Plant viability and fungal identification

None of the treatments had a negative influence on callus or initial shoot growth. The viability of planting material was estimated to be of 94% and 92% for the 2-year-old and 3-year-old plants at the end of growing season, respectively. After HWT, six and four out of the 50 grafted plants stock tested positive for *Diplodia seriata* and *Neofusicoccum parvum*, respectively, other fungi associated with GTDs. No black-foot and Petri disease pathogens were isolated from hot-water treated plants. In the 2-year-old plants, a total of 1,650 Petri disease (83.6% from the basal end of the rootstock and 16.4% from roots) and 896 black-foot disease pathogens (15.8% from the basal end of the rootstock and 84.2% from roots) isolates were collected. Petri disease pathogens were identified as *C. luteo-olivacea* (57.8%), followed by *Pa. chlamydospora* (27.3%) and *Pm. minimum* (14.9%).

Black-foot pathogens were identified as *Dactylonectria torresensis* (66.4%), followed by *Dactylonectria macrodidyma* (22.6%), *Ilyonectria liriodendri* (6.2%) and *Dactylonectria alcacerensis* (4.8%). In the 3-year-old plants, a total of 1,825 Petri disease (89.4% from the basal end of the rootstock and 10.6% from roots) and 1,632 black-foot pathogens (26.9% from the basal end of the rootstock and 73.1% from roots) isolates were collected. Petri disease pathogens were identified as *C. luteo-olivacea* (54.6%), followed by *Pa. chlamydospora* (31.1%) and *Pm. minimum* (14.3%). Black-foot pathogens were identified as *D. torresensis* (66.0%), followed by *I. liriodendri* (16.0%), *D. macrodidyma* (9.2%), *Ilyonectria robusta* (4.4%), *D. alcacerensis* (2.5%) and *Ilyonectria pseudodestructans* (1.8%). Representative black-foot and Petri diseases isolate sequences obtained in this study were deposited to GenBank (Supplementary Table S1).

*Trichoderma atroviride* was isolated from 30 and 22% of the 2-year-old and 3-year-old plants, respectively. *Trichoderma koningii* was isolated from 12 and 18% of the 2-year-old and 3-year-old plants, respectively. Our attempts to isolate *P. oligandrum* were unsuccessful.

### 3.2 Disease incidence and disease severity in grafted plants

Neither field site, nor block, nor its interaction significantly affected the DI and DS (*P*>0.05, ANOVA not shown). Therefore, data from both field sites were combined and analysed together. There was a significant effect of treatment on mean Petri disease incidence values in the roots and the basal ends for both 2-year-old and 3-year-old plants (Table 1). In the 3-year-old plants, percentage of infected plants (DI) in the basal ends were significantly lower in treatments with *T. atroviride* SC1 (T3) (40.2% ± 8.3) than in the control treatment (61.5% ± 5.6) (Figure 1A). In both 2-year-old and 3-year-old plants, percentage of infected plants (DI) in the roots were significantly lower in treatments with *P. oligandrum* Po37 (T5) (2-year-old plants: 7.5% ± 1.4, and 3-year-old plants: 4.8% ± 1.3) than in the control treatment (2-year-old plants: 23.1% ± 2.8, and 3-year-old plants: 18.3% ± 3.9) (Figure 1B). Biocontrol treatments had a significant effect on mean Petri disease severity in basal ends of 2-year-old plants, and in roots and basal ends for 3-year-old plants (Table 1). *T. atroviride* SC1 (T3) in the 2-year-old plants (19.4% ± 1.4) and both *Trichoderma* spp. treatments (T2: 25.5% ± 2.5, and T3: 25.8% ± 2.3) in the 3-year-old plants significantly reduced the percentage of DS in the basal ends compared to the control treatment (2-year-old plants: 36.1% ± 4.3, and 3-year-old plants: 39.5% ± 4.9) (Figure 1A). *Trichoderma* spp. treatments (T2: 9.1% ± 1.3, and T3: 10.8% ± 1.8) resulted in significant lower DS in roots of the 3-year-old plants than the control treatment (16.8% ± 3.8) (Figure 1B).

**Table 1.** Effects of variables on disease incidence (DI) and disease severity (DS) in roots and basal ends of rootstock of grapevine grafted plants.

| Petri disease | 2-year-old plants |  |  |  |  | 3-year-old plants |  |  |  |  |
| --- | --- | --- | --- | --- | --- | --- | --- | --- | --- | --- |
|  | DI (%) |  |  | DS (%) |  | DI (%) |  |  | DS (%) |  |
|  | df<sup>a</sup> | MS<sup>b</sup> | *P* value<sup>c</sup> | MS | *P* value | df | MS | *P* value | MS | *P* value |
| Roots |  |  |  |  |  |  |  |  |  |  |
| Block | 3 | 0.0034 | 0.9920 | 0.0129 | 0.8115 | 3 | 0.0036 | 0.5770 | 0.0332 | 0.9112 |
| Treatment | 5 | 0.0162 | 0.0003 | 0.0196 | 0.2107 | 5 | 0.0202 | 0.0010 | 0.0708 | 0.0418 |
| Error | 63 | 0.0029 |  | 0.0133 |  | 63 | 0.0042 |  | 2.1901 |  |
| Basal ends |  |  |  |  |  |  |  |  |  |  |
| Block | 3 | 0.03102 | 0.7304 | 0.01033 | 0.9031 | 3 | 0.03008 | 0.4951 | 0.02974 | 0.9954 |
| Treatment | 5 | 0.03810 | 0.0411 | 0.00806 | 0.0397 | 5 | 0.03400 | 0.0433 | 0.01761 | 0.0462 |
| Error | 63 | 0.01686 |  | 0.01890 |  | 63 | 0.01770 |  | 0.02023 |  |
| Black-foot disease | 2-year-old plants |  |  |  |  | 3-year-old plants |  |  |  |  |
|  | DI (%) |  |  | DS (%) |  | DI (%) |  |  | DS (%) |  |
|  | df | MS | *P* value | MS | *P* value | df | MS | *P* value | MS | *P* value |
| Roots |  |  |  |  |  |  |  |  |  |  |
| Block | 3 | 0.0141 | 0.9945 | 0.0071 | 0.9110 | 3 | 0.1212 | 0.9932 | 0.0448 | 0.8234 |
| Treatment | 5 | 0.0024 | 0.9973 | 0.0121 | 0.7817 | 5 | 0.0082 | 0.9162 | 0.0044 | 0.0526 |
| Error | 63 | 0.0393 |  | 0.0247 |  | 63 | 0.0282 |  | 0.0467 |  |
| Basal ends |  |  |  |  |  |  |  |  |  |  |
| Block | 3 | 0.0076 | 0.8712 | 0.0081 | 0.7883 | 3 | 0.0135 | 0.9003 | 0.0328 | 0.9991 |
| Treatment | 5 | 0.0282 | 0.0413 | 0.0328 | 0.0417 | 5 | 0.0167 | 0.0389 | 0.0151 | 0.9404 |
| Error | 63 | 0.0136 |  | 0.0501 |  | 63 | 0.0233 |  | 0.0613 |  |

<sup>a</sup> df = degrees of freedom.<sup>b</sup> MS = mean square.<sup>c</sup> Significance level $P < 0.05$ .

Analysis of variance showed no significant effect of biocontrol treatments on black-foot disease incidence and severity in roots of both 2-year-old and 3-year-old plants (Table 1). There was a significant effect of treatment on mean black-foot disease incidence values in the basal ends for both 2-year-old and 3-year-old plants (Table 1). In the 2-year-old plants, all treatments resulted in significant lower DI in the basal ends than the control treatment (Figure 2A). In the 3-year-old plants, percentage of infected plants (DI) in the basal ends were significantly lower in treatments with *Streptomyces* sp. E1 + R4 (T1) (4.8% ± 2.1) than in the control treatment (13.2% ± 3.2) (Figure 2A). There was a significant effect of treatment on mean black-foot disease severity values in the basal ends of 2-year-old plants (Table 1). *Streptomyces* sp. E1 + R4 (T1) (10.0% ± 2.3) significantly reduced the percentage of DS in the basal ends compared to the control treatment (19.1 ± 0.8) (Figure 2A).

### 3.3 Fungal species incidence and severity in grafted plants

Considering the fungal species within each disease individually, *P. oligandrum* Po37 (T5) and *T. atroviride* SC1 (T3) significantly reduced the DI of *C. luteo-olivacea* in the roots and the basal ends, respectively, of 2-year-old plants compared to the control treatment (*P*<0.05, ANOVA not shown) (Table 2). Percentage of DI in the roots of both 2-year-old and 3-year-old plants, and DS in the roots of 2-year-old plants caused by *Pa. chlamydospora* were significantly lower in treatments with *P. oligandrum* Po37 (T5) than in the control treatment (*P*<0.05, ANOVA not shown) (Table 2). In the 3-year-old plants, *T. atroviride* SC1 (T3) significantly reduced both DI and DS caused by *Pa. chlamydospora* in the basal ends compared to the control treatment (*P*<0.05, ANOVA not shown) (Table 2). Both *T. koningii* TK7 (T2) and *P. fluorescens + B. atrophaeus* (T4) treatments resulted in significant lower DI caused by *Pm. minimum* in the roots of 2-year-old plants than the control treatment (*P*<0.05, ANOVA not shown) (Table 2). Furthermore, *T. koningii* TK7 (T2) treatment resulted in significant lower DS caused by *Pm. minimum* in the roots of 3-year-old plants than the control treatment (*P*<0.05, ANOVA not shown) (Table 2). *T. atroviride* SC1 (T3) significantly reduced the DS of *Pm. minimum* in the roots of both 2-year-old and 3-year-old plants compared to the control treatment (*P*<0.05, ANOVA not shown) (Table 2).

**Table 2.** Mean disease incidence (DI) and severity (DS) of Petri disease pathogens isolated from the roots and basal ends of the rootstock of 2-year-old and 3-year-old grafted plants subjected to various treatments prior to planting in two fields in Logroño (La Rioja).

| *Cadophora luteo-olivacea* |  |  |  |  |  |  |  |  |
| --- | --- | --- | --- | --- | --- | --- | --- | --- |
|  | 2-year-old plants |  |  |  | 3-year-old plants |  |  |  |
|  | DI (%)<sup>a</sup> |  | DS (%)<sup>a</sup> |  | DI (%) |  | DS (%) |  |
|  | Roots | Basal ends | Roots | Basal ends | Roots | Basal ends | Roots | Basal ends |
| T1. *Streptomyces* sp. E1 + R4 | 8.1 a | 24.4 ab | 20.4 a | 26.7 a | 6.9 a | 25.6 a | 19.4 a | 31.3 a |
| T2. *Trichoderma koningii* TK7 | 5.6 ab | 34.4 a | 20.4 a | 24.5 a | 5.6 a | 34.4 a | 19.6 a | 28.4 a |
| T3. *Trichoderma atroviride* SC1 | 6.9 ab | 20.6 b | 21.0 a | 30.3 a | 5.0 a | 28.1 a | 19.2 a | 27.6 a |
| T4. *Pseudomonas fluorescens* + *Bacillus atrophaeus* | 6.3 ab | 31.3 ab | 22.3 a | 30.7 a | 6.3 a | 31.3 a | 20.4 a | 35.0 a |
| T5. *Pythium oligandrum* Po37 | 3.8 b | 31.9 ab | 21.3 a | 30.3 a | 5.0 a | 31.9 a | 17.5 a | 31.0 a |
| Control (C) | 7.5 a | 38.1 a | 25.2 a | 33.8 a | 7.5 a | 36.9 a | 22.3 a | 33.4 a |
| LSD (*P* = 0.05) | 2.7 | 8.2 | 7.5 | 7.7 | 2.5 | 7.8 | 7.0 | 7.5 |
| *Phaeomoniella chlamydospora* |  |  |  |  |  |  |  |  |
|  | 2-year-old plants |  |  |  | 3-year-old plants |  |  |  |
|  | DI (%) |  | DS (%) |  | DI (%) |  | DS (%) |  |
|  | Roots | Basal ends | Roots | Basal ends | Roots | Basal ends | Roots | Basal ends |
| T1. *Streptomyces* sp. E1 + R4 | 6.3 a | 11.3 a | 10.0 ab | 27.6 a | 7.5 a | 15.6 a | 9.2 a | 28.2 ab |
| T2. *Trichoderma koningii* TK7 | 6.3 a | 18.8 a | 10.6 ab | 25.0 a | 8.1 a | 18.8 a | 14.2 a | 35.0 a |
| T3. *Trichoderma atroviride* SC1 | 7.5 a | 14.4 a | 14.8 ab | 29.8 a | 8.1 a | 6.3 b | 15.8 a | 15.0 b |
| T4. *Pseudomonas fluorescens* + *Bacillus atrophaeus* | 6.3 a | 17.5 a | 13.3 ab | 28.5 a | 7.5 a | 17.5 a | 11.7 a | 38.7 a |
| T5. *Pythium oligandrum* Po37 | 1.9 b | 22.5 a | 7.5 b | 20.9 a | 0.6 b | 22.5 a | 12.5 a | 31.0 a |
| Control (C) | 9.4 a | 20.6 a | 15.4 a | 28.3 a | 9.4 a | 20.6 a | 10.6 a | 34.6 a |
| LSD (*P* = 0.05) | 3.1 | 5.5 | 3.6 | 7.4 | 3.3 | 3.8 | 3.8 | 7.1 |
| *Phaeoacremonium minimum* |  |  |  |  |  |  |  |  |
|  | 2-year-old plants |  |  |  | 3-year-old plants |  |  |  |
|  | DI (%) |  | DS (%) |  | DI (%) |  | DS (%) |  |
|  | Roots | Basal ends | Roots | Basal ends | Roots | Basal ends | Roots | Basal ends |
| T1. *Streptomyces* sp. E1 + R4 | 4.4 ab | 11.3 a | 16.3 a | 15.1 a | 4.0 a | 15.0 a | 13.8 a | 21.0 a |
| T2. *Trichoderma koningii* TK7 | 3.1 b | 16.9 a | 18.8 a | 13.6 a | 4.1 a | 16.5 a | 2.5 b | 9.5 b |
| T3. *Trichoderma atroviride* SC1 | 5.0 ab | 11.3 a | 7.1 b | 11.8 a | 3.9 a | 14.7 a | 3.8 b | 22.4 a |
| T4. *Pseudomonas fluorescens* + *Bacillus atrophaeus* | 3.1 b | 17.5 a | 23.8 a | 12.8 a | 4.3 a | 15.1 a | 15.0 a | 17.7 ab |
| T5. *Pythium oligandrum* Po37 | 3.8 ab | 16.9 a | 16.3 a | 10.3 a | 4.0 a | 15.8 a | 12.5 a | 18.3 ab |
| Control (C) | 5.6 a | 16.9 a | 21.7 a | 17.0 a | 4.0 a | 17.2 a | 12.5 a | 19.9 ab |
| LSD (*P* = 0.05) | 2.3 | 3.6 | 3.5 | 3.3 | 2.7 | 3.1 | 3.6 | 3.5 |

<sup>a</sup> At each plant part, percentages of disease incidence (DI) and disease severity (DS) are the mean of 160 plants analyzed (40 plants per replicate). Values in the same column followed by the same letter do not differ significantly (*P*=0.05).

Regarding black-foot pathogens, all treatments significantly reduced the DI of *D. torresensis* and *D. macrodidyma* in the basal ends of 2-year-old plants compared to the control treatment (*P*<0.05, ANOVA not shown) (Table 3). *Streptomyces* sp. E1 + R4 (T1) significantly reduced *D. torresensis* DS in the basal ends of 2-year-old plants and DI in the basal ends of 3-year-old plants compared to the control treatment (*P*<0.05, ANOVA not shown) (Table 3). In both the 2-year-old and 3-year-old plants, percentages of DI in the roots and DS in the basal ends caused by *D. macrodidyma* were significantly lower in treatments with *Streptomyces* sp. E1 + R4 (T1) than in the control treatment (*P*<0.05, ANOVA not shown) (Table 3). *T. atroviride* SC1 (T3) also resulted in significant lower DS in the basal ends of 3-year-old plants than the control treatment (*P*<0.05, ANOVA not shown) (Table 3). Low levels of *Trichoderma* spp. (<30%) were isolated from roots and basal ends of 2-year-old and 3-year-old plants subjected to T2 and T3 treatments in both fields.

**Table 3.**
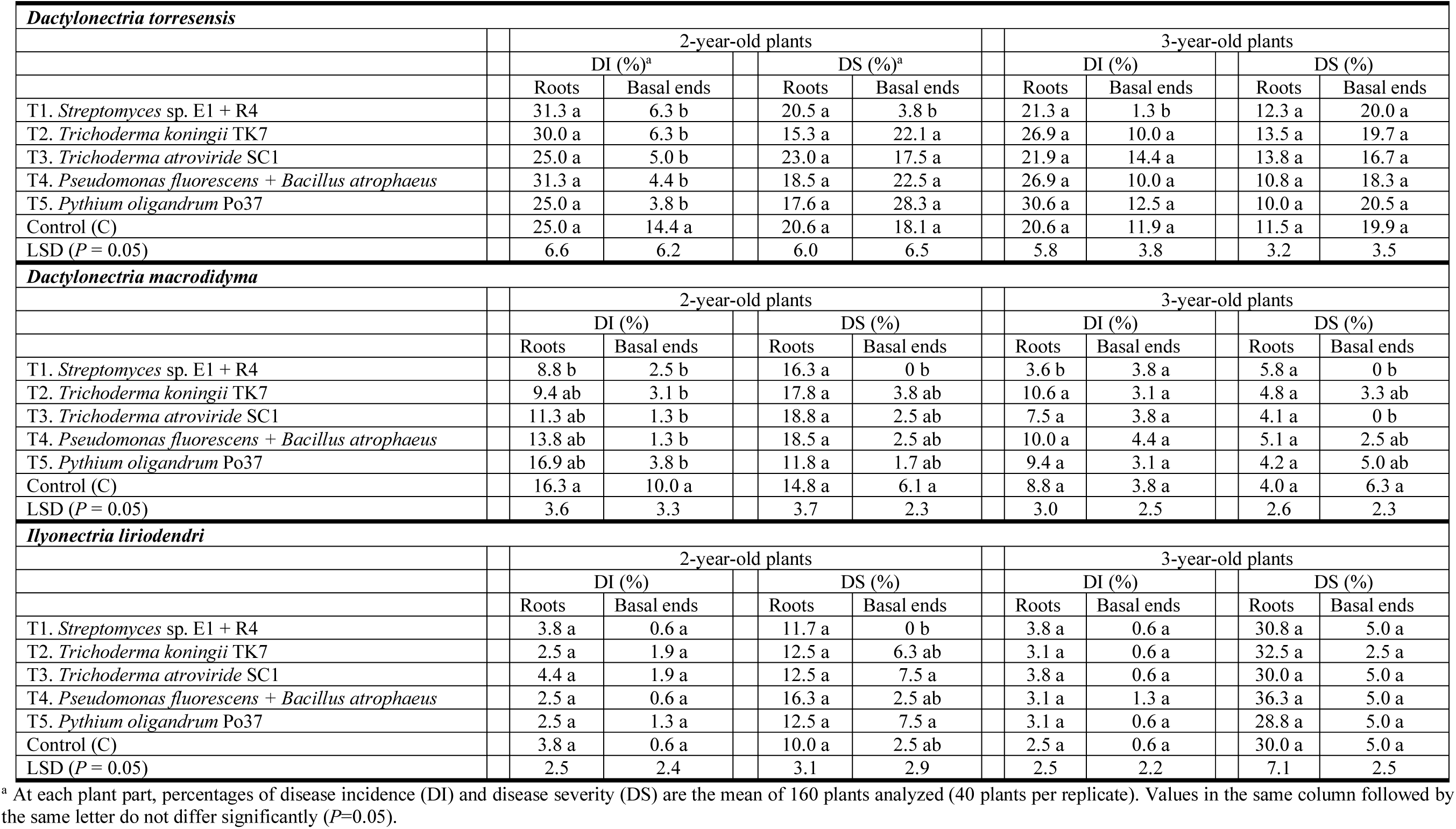
Mean disease incidence (DI) and severity (DS) of the most prevalent black-foot disease pathogens isolated from the roots and basal ends of rootstock of 2-year-old and 3-year-old grafted plants subjected to various treatments prior to planting in two fields in Logroño (La Rioja).

The percentage of reduction (PR) was calculated for treatments statistically different from the control in the 3-years-old plants (Table 4). In roots, *P. oligandrum* Po37 (T5) provided 93.6% disease incidence reduction of *Pa. chlamydopora*. On *Trichoderma* spp. treated plants, there was a reduction in *Pm. minimum* severity when compared with untreated controls, which ranged from 80% for *T. koningii* TK7 (T2) and 69.6% for *T. atroviride* SC1 (T3). In the basal ends, *T. atroviride* SC1 (T3) provided 69.4% disease incidence and 56.6% disease severity reduction of *Pa. chlamydopora*, while *T. koningii* TK7 (T2) provided 52.3% disease severity reduction of *Pm. minimum.* None of the BCA treatments statistically reduced the disease incidence and severity of black-foot disease fungi in roots (Tables 3 and 4). In the basal ends, *Streptomyces* sp. E1 + R4 (T1) reduced the incidence of *D. torresensis* and the severity of *D. macrodidyma* by 89.1 and 100%, respectively. *T. atroviride* SC1 (T3) provided 100% disease severity reduction of *D. macrodidyma*.

**Table 4.**
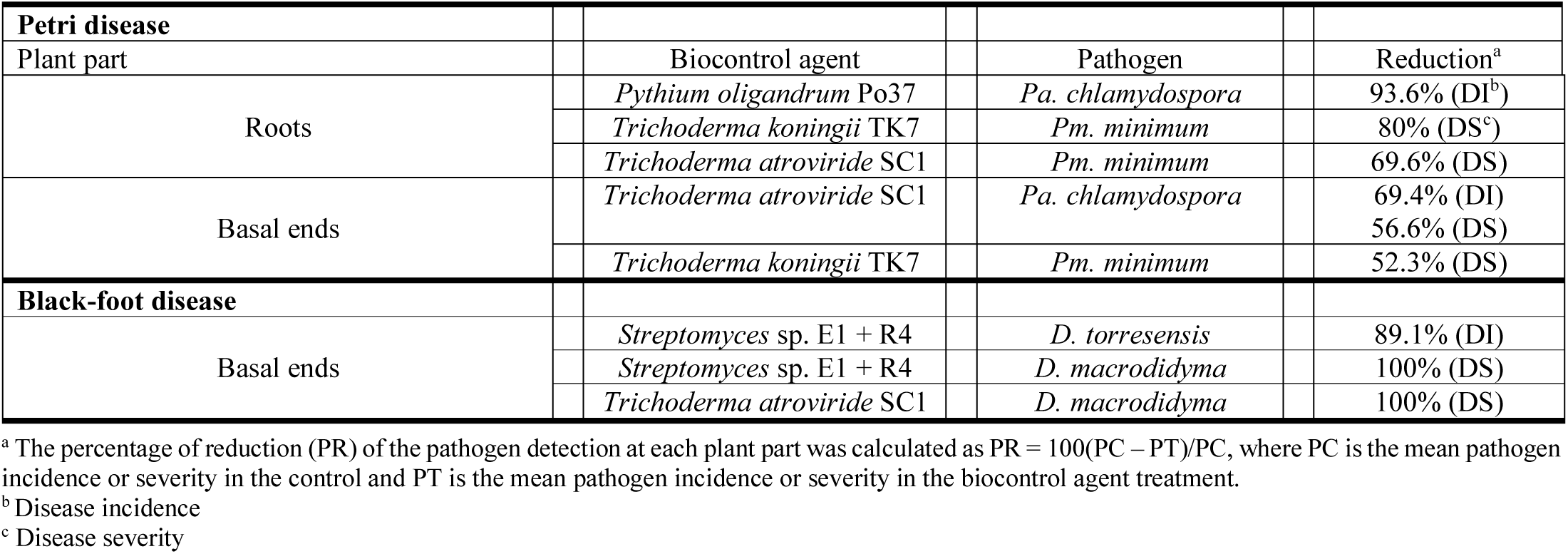
Pathogen reduction achieved by BCA treatments in the 3-year-old plants, associated with Petri and black-foot disease.

### 3.4 Root and shoot weights in grafted plants, and physicochemical properties of the soil

Analysis of variance showed no significant effect of biocontrol treatments on the shoot weight of 3-year-old plants (*P*>0.05, ANOVA not shown) (Figure 3). Mean shoot weight ranged from 55.3 g ± 5.7 (T3) to 64.9 g ± 8.2 (T2). Biological control treatments had a significant effect on the root weight of 3-year-old plants (*P*<0.05, ANOVA not shown) (Figure 3). Mean root weight ranged from 41.9 g ± 3.7 (T3) to 52.9 g ± 2.9 (C). All treatments resulted in significant lower root weight than the control treatment (Figure 3). Analyses of variance indicated no significant differences for the soil physicochemical properties between fields (*P*>0.05, ANOVA not shown).

## 4 DISCUSSION

This study represents the first approach to evaluate the effectiveness of different antagonistic microorganisms (bacteria, fungi and an oomycete) applied preventively to control black-foot and Petri diseases under field conditions. The use of BCA against soilborne pathogens are on the forefront of research; however, most experiences are on a laboratory scale, thus avoiding the problems related to the production of large quantities of antagonists and their formulations, and disease control trials are performed in simplified environment such as growth chambers or experimental greenhouses, thus avoiding the risk of large-scale experiments in the field.

In this study, Petri disease infection was mainly detected in the basal ends of the rootstock, while fungi associated with black-foot disease were most frequently isolated from roots. *D. torresensis* was the most frequent isolated species (>60%) associated with black-foot disease at both plant ages. This agrees with previous research carried out on black-foot in Europe.^9-11^ Regarding Petri disease, more than 80% of the fungi were identified as *C. luteo-olivacea* and *Pa. chlamydospora* at both plant ages. Both fungal species were frequently isolated from nursery stock and young vines worldwide.^3^

In our specific pathosystems, the effectiveness of some BCA in reducing the incidence and severity of both diseases under field conditions were dependent on the plant part analysed and the plant age. *Streptomyces* sp. E1 + R4 treatment was highly effective in reducing black-foot disease incidence at both plants ages and the severity of 2-year-old plants in the basal ends. However, the effect of these actinobacteria against Petri disease pathogens after 2 years in the field was very low. In contrast, Álvarez-Pérez *et al*.^38^ evaluated the effectiveness of these bacterial strains individually, previously isolated from the endo- (strain E1) and rhizosphere (strain R4) of the grapevine root system, for black-foot and Petri diseases control in 1-year-old grafted plants under field conditions by partially immersing the grafts (up to 10 cm depth) in a rooting hormone solution containing the actinobacteria for 24 h at room temperature. They found significant reductions of the infection rates at the lower end of the rootstock of the fungal pathogens *Dactylonectria* sp., *Ilyonectria* sp., *Pm. minimum* and *Pa. chlamydospora*.^38^ These differences in the effectiveness of the bacteria against Petri disease between experiments could be due to the commonly unpredictably behaviour of BCA when tested in different environments.^58^

Other bacterial treatment tested in our study was a commercial product containing *Pseudomonas fluorescens* and *Bacillus atrophaeus*. No biocontrol effect of this treatments was observed on fungal pathogens associated with black-foot and Petri diseases. Despite this fact, some strains of these bacterial species have been previously reported as plant growth-promoting bacteria (PGPB) and have been found to be potential BCA of plant diseases in several crops.^59-62^ In grapevine, different *P. fluorescens* strains were identified as prospective new BCA against *Botrytis cinerea*^63^ and to induce systemic resistance against *Plasmopara viticola* and *B. cinerea* by priming common and distinct defensive pathways.^64^

The *in vitro* effects of beneficial bacteria in reducing GTDs has been also tested.^65,66^ *Bacillus subtilis* AG1 showed promising in reducing the growth of *Lasiodiplodia theobromae, Pa. chlamydospora*, and *Pm. minimum* in an artificial culture medium.^34^ Rezgui *et al*.^37^ recently identified antagonistic traits against GTDs pathogens of several *B. subtilis* strains inhabiting the wood tissues of mature grapevines in Tunisia. The antagonistic activity of *Pantoea agglomerans* and *Bacillus pumilus* against *N. parvum* and *Pa. chlamydospora* was demonstrated in inoculated ungrafted grapevine cuttings.^35,36^ A recent study performed by Trotel-Aziz *et al*.^40^ highlighted the effect of *B. subtilis* strain PTA-271 to efficiently attenuate the characteristic Botryosphaeria dieback symptoms caused by *N. parvum*.

Most studies on biological control of GTDs have examined the application of *Trichoderma* spp. in grapevine nurseries and young vineyards.^21-25,29^ In our study, we individually evaluated two *Trichoderma*-based products containing *T. koningii* strain TK7 and *T. atroviride* strain SC1. A certain effect was observed in reducing *Pm. minimum* disease incidence for 2-year-old plants and disease severity for 3-year-old plants at the root level by *T. koningii* TK7 treatment. Little information is still available related to the biocontrol effect of TK7 strain to combat plants’ fungal pathogens. Howell *et al*.^67^ showed that the application of *T. koningii* TK7 to cotton seeds before planting was ineffective to control cotton seedlings damping-off in artificially *Rhizoctonia solani*-infested cotton field soil flats. Other strains of this species have been recently reported as potential BCA for fungal pathogens in different crops, such as *Fusarium oxysporum* f. sp. *melonis* in melon^68^ or *Sclerotium rolfsii* in groundnut.^69^

*Trichoderma atroviride* SC1 was effective in reducing *Pa. chlamydospora* disease incidence and severity in the basal ends of 3-year-old plants. In accordance with our results, a study carried out in Spain by Berbegal *et al*.^33^ also found reductions in the incidence and severity of *Pa. chlamydospora* and *Pm. minimum* when analysed the rootstock basal end and root system of 1-day *T. atroviride* SC1 inoculated grafted plants in nurseries. In the same study, Berbegal *et al*.^33^ also evaluated the effect of *T. atroviride* SC1 treatment in two fields during two growing seasons. The basal parts of the treated plants were soaked for 1 h in a *T. atroviride* SC1 suspension before planting, observing no BCA effect on incidence and severity of black-foot disease associated pathogens and significant reductions on pathogens associated with Petri disease at both fields after the first growing season. In Italian grapevine nurseries, the application of *T. atroviride* strain SC1 at several stages of the nursery process (pre-storage and pre-grafting hydration, stratification, callusing, and rooting) protected plants from infection by *Pm. minimum* and *Pa. chlamydospora* after a single artificial inoculation with both pathogens following the grafting stage.^28^

Regarding *P. oligandrum* Po37 treatment, a significant reduction of Petri disease incidence and severity was observed in 2-year-old plants and disease incidence in 3-year-old plants, at roots level. Yacoub *et al*.^41^ reported a significant reduction in necrosis length caused by *Pa. chlamydospora* when the roots of ‘Cabernet Sauvignon’ cuttings were colonized by different *P. oligandrum* strains. The efficacy of *P. oligandrum* strain Sto7 in reducing the necrosis length caused by *N. parvum* and *Pa. chlamydospora* was demonstrated on grafted young ‘Cabernet Sauvignon’ vines cultivated in a nursery greenhouse, separately or in combination with two bacterial strains previously isolated from vineyards.^42^ The ability of *P. oligandrum* strain Po37 to act as an inducer of plant systemic resistance against pathogens is thought to be due to the presence of three elicitin-like proteins in its genome.^70^

Diverse formulations (dry or water suspensions), application methods and times of exposure of plants to BCA have been tested in the different studies carried out to assess the biocontrol potential of antagonist microorganisms.^28,30,33,38,41,42,71-73^ In our assay, a 24-h soaking of the trimmed root systems and the basal end of the plants in BCA water suspensions was carried out before planting, but the percentage of *Trichoderma* spp. recovery was low in all cases (<30%). In this sense, Halleen *et al*.^74^ were also able to only isolate a 2.3% of *Trichoderma* spp. from the basal ends of the rootstock and none from roots of grafted plants subjected to *Trichoderma* treatments, applied by dipping the basal ends of the rootstock for 1 min before planting, after 7 months in a nursery field. In a recent study, González-García *et al*.^72^ evaluated the colonization efficiency of *Streptomyces* sp. in the root system by comparing two inoculation methods, plant immersion in a bacterial suspension or direct injection of the bacterial suspension into the vegetal tissues and concluded that both methods allowed effective BCA colonization. This is also in accordance with Berbegal *et al*.^33^ who used 24-h soak in *T. atroviride* SC1 water suspension to inoculate 110R rootstock cuttings before grafting, with percentages of recovery over 80% at both nursery and vineyard experiments. Van Jaarsveld *et al*.^73^ evaluated different methods of application of *T. atroviride* on commercially planted nursery vines and concluded that dipping of basal ends in the *Trichoderma* dry formulation consistently gave higher colonization percentages than the 1-h soak of bases of vines before planting or *Trichoderma* field drenching. Further research is needed to evaluate the effectiveness of soaking vines in *T. koningii* TK7 or *T. atroviride* SC1 dry formulations compared to soaking vines for 24-h in BCA water suspensions before planting.

Biological control agent treatments did not affect the shoot weight, and root weight was significantly lower for all BCA treatments with respect to the untreated control at the end of the second growing season (3-year-old plants). The impact of BCA treatments on grapevine development was very variable on previous research.^29,73,74^ *Trichoderma* spp. and *B. subtilis*-based treatments resulted in lower mean root and shoot dry weight values when compared with the negative controls.^29^ Nevertheless, Halleen *et al*.^74^ found that none of the *Trichoderma* formulations tested yielded plants with roots or shoots mass significantly different than the water treated controls. Berbegal *et al*.^33^ observed a significantly higher undried shoot weight for *T. atroviride* SC1 treated plants at the end of the first growing season, but this effect was not observed in the second growing season. Likewise, the application of actinobacteria to grafted grapevine plants did not show a significant effect, either positive or negative, on plants growth.^38^ In contrast, Fourie *et al*.^22^ observed that *T. harzianum* treatments significantly improved root development but not shoot mass in comparison with the control vines in nurseries. Several studies indicate that BCA treatments can enhance the growth of other crops, such as tomato^75^ or rice.^76^ All this variability could be related to the lack of proper long-standing implantation by these antagonist microorganisms in grapevine roots or the vigour level of the rootstock cultivar tested. BCA are living organisms whose activities depend mainly on the different physicochemical environmental conditions to which they are subjected,^77^ and the greatest long-term effects probably occur with rhizosphere-competent strains with the ability to colonize and grow in association with plant roots.^78^

## 5 CONCLUSIONS

This study highlighted the potential of some BCA to reduce GTD infection under field conditions. No single BCA application was able to control both diseases. Further studies should evaluate the combination of the disease-suppressive activity of two or more beneficial microbiomes in a biocontrol preparation against black-foot and Petri diseases. Our results also open up the possibility to combine the application of BCA as a pre-planting strategy with other measures in an Integrated Pest Management (IPM) programme against GTDs. For example, BCA can be applied after hot-water treatment (HWT) of dormant grafted plants or after soil biofumigation. In this regard, recent research highlighted the effectiveness of HWT at 53°C for 30 min^45^ and white mustard biofumigation^30^ to reduce GTD incidence in planting material and grapevine nursery soil, respectively.

## ACKNOWLEDGEMENTS

We thank researchers of the “Instituto de Investigación de la Viña y el Vino” in the “Universidad de León” for providing the *Streptomyces* sp. strains. M.P. Martínez-Diz was supported by the FPI-INIA program from the INIA. D. Gramaje was supported by the Ramón y Cajal program, Spanish Government (RYC-2017-23098).

